# Three dimensional two-photon imaging of neuronal activity in freely moving mice using a miniature fiber coupled microscope with active axial-scanning

**DOI:** 10.1101/226431

**Authors:** Baris N. Ozbay, Gregory L. Futia, Ming Ma, Victor M. Bright, Juliet T. Gopinath, Ethan G. Hughes, Diego Restrepo, Emily A. Gibson

## Abstract

We present a miniature head mounted two-photon fiber-coupled microscope (2P-FCM) for neuronal imaging with active axial focusing enabled using a miniature electrowetting lens. Full three-dimensional two-photon imaging of GCaMP6s showing individual neuron activity in multiple focal planes was achieved in a freely-moving mouse. Two-color simultaneous imaging of GFP and tdTomato fluorescence is also demonstrated. Additionally, dynamic control of the axial scanning of the electrowetting lens allows tilting of the focal plane enabling cells in multiple focal planes to be imaged simultaneously. Two-photon imaging allows increased penetration depth in tissue yielding a working distance of 450 μm with an additional 180 μm of active axial focusing. The objective NA is 0.45 with a lateral resolution of 1.8 μm, an axial resolution of 10 μm, and a field-of-view of 240 μm diameter. The 2P-FCM has a weight of only ∼2.5 g and is capable of repeatable and stable head-attachment. The 2P-FCM with dynamic axial scanning provides a new capability to record from functionally distinct neuronal layers, opening new opportunities in neuroscience research.

## Introduction

Two-photon excitation (2PE) microscopy^1,2^ has become a fundamental tool for probing *in vivo* neuronal circuits because it yields optical access deeper into tissue compared to single-photon fluorescence microscopy. 2PE microscopy allows for optical sectioning while imaging in scattering tissue and, when used in combination with rapid axial-scanning, allows for the retrieval of three-dimensional (3D) representations of neuronal structure and activity^3,4^. As optical proteins, such as genetically-encoded fluorescence Ca^2+^ sensors, have continued to improve^5^, 2PE microscopy has been a dominant tool for *in vivo* imaging in neuroscience research.

In the previous decade, there has been a technological push to allow for imaging of neuronal activity of freely-moving animals to study neuronal circuitry in behaviors such as spatial navigation^6^, social behavior^7^, or prey capture^8^. Several commercial devices have been developed for head mounted wide-field epifluorescence imaging in freely-behaving mice^9,10^. Furthermore, a variety of miniature laser-scanning fiber-coupled microscopes (FCMs) for confocal and 2PE imaging have been developed^11-13^. However, all of these devices suffer from lack of active axial-scanning that greatly limits their applications. In order to fully study the spatially complex neuronal interconnections using miniaturized laser-scanning microscopy, it is required to image in 3D.

Lateral imaging with a FCM is accomplished by a variety of techniques that scan the excitation laser focal spot in two axes at the sample. Axial-scanning requires an additional third degree-of-freedom that can complicate the design of the optics and mechanics of the FCM. Although several axial-scanning solutions have been demonstrated for FCMs, these have not been sucessfully translated to *in vivo* neuronal recording in freely-moving animals due to various drawbacks in these methods. For example, MEMS mirrors can be modified to include an axial scanner, but this adds mechanical and optical complexity that limit miniaturizion^14^. Axial-scanning can also be accomplished using an electrically tunable lens (ETL) to adjust the focal length (FL) of the imaging system. ETLs are attractive because they are transmissive and can be placed directly in the optical path, so they don’t require the extra complexity of reflective surfaces in the optical path. ETLs can be enabled by shape-memory alloys^15^ or pneumatic actuation^16^, however, these solutions are too slow for fast 3D scanning and not easily miniaturized. Other types of ETLs include shape-changing polymer lenses^17,18^, pressure-driven lenses^19^, and responsive hydrogel lenses^20^. However, these suffer from not being adaptable to miniaturization, large optical aberrations, and/or being too susceptible to motion and orientation for freely-behaving animal imaging. Electrowetting tunable lens (EWTL) technology is an ideal candidate for axial-scanning that can be integrated into FCMs. EWTLs can be miniaturized to mm-scales, are unaffected by motion and orientation, and can have very low power requirements (< 20 milliWatts)^21-23^.

In this article, we describe the use of EWTL technology to develop the first two-photon fiber coupled microscope (2P-FCM) capable of performing active axial-scanning for imaging in an awake and freely-behaving mouse. We demonstrate the 2P-FCM for *in vivo* Ca^2+^-imaging of neurons in the cortex. Ca^2+^-transients are observed from distinct focal planes. We show the ability to tilt the focal plane during imaging using the EWTL, allowing imaging of cellular populations in planes other than those perpendicular to the optical axis. Additionally, the 2P-FCM can perform simultaneous two-color imaging, demonstrated by imaging GFP and tdTomato, showing the ability to measure from distinct cell types. The 2P-FCM enclosure is lightweight and can be repeatedly attached and removed from the animal with stable imaging over the same brain region on separate days. The imaging field-of-view (FOV) is a cylindrical volume with 240 μm diameter x 180 μm depth. The 2P-FCM presented here is, to the best of our knowledge, the first device to attain 3D-imaging of neuronal activity in the brain of a freely-moving mouse.

## Results

### Two-photon fiber-coupled microscope design

The overall setup is shown in **Figure 1A**. A 1.0 meter long flexible coherent fiber bundle (CFB) is used as an optical relay to couple the head-mounted miniature optics to a bench-top laser scanning two-photon microscope (2PM). The lateral scanning of the excitation laser is done at the proximal end of the CFB using the galvanometric scanner in the 2PM to translate the lateral position of the laser focus through the spatially coherent fiber-cores from the proximal to the distal end. This lateral scanning technique has been demonstrated previously for both confocal and 2PE imaging and avoids the need for miniaturized scanners at the distal end^24-26^, allowing the head-mounted design to be lighter weight and contain fewer complex components. The tradeoff is the lower resolution, which is limited by the number and spacing of the fiber cores. The surface of the CFB is shown in **Figure 1B**, which has an outer diameter of 650 µm, and an effective imaging diameter of 550 µm. The CFB has ∼15,000 individual fiber cores, with a core diameter of ∼2.9 µm and core-to-core spacing of ∼4.5 µm. In order to maintain the ultrashort pulse-durations required for efficient 2PE, the laser pulse must be pre-compensated to account for nonlinear spectral narrowing and chromatic dispersion that arise from propagating high intensity light through the CFB cores. We implemented previously described pre-compensation techniques to recover a ∼100 fs pulse width at the output of the CFB. These were implemented using a polarization-maintaing single-mode fiber (PMF) and grating-pair based pulse stretcher to spectrally broaden and apply negative chirp to the pulse, respectively (see *Methods* section for more details).

**Figure 1.**
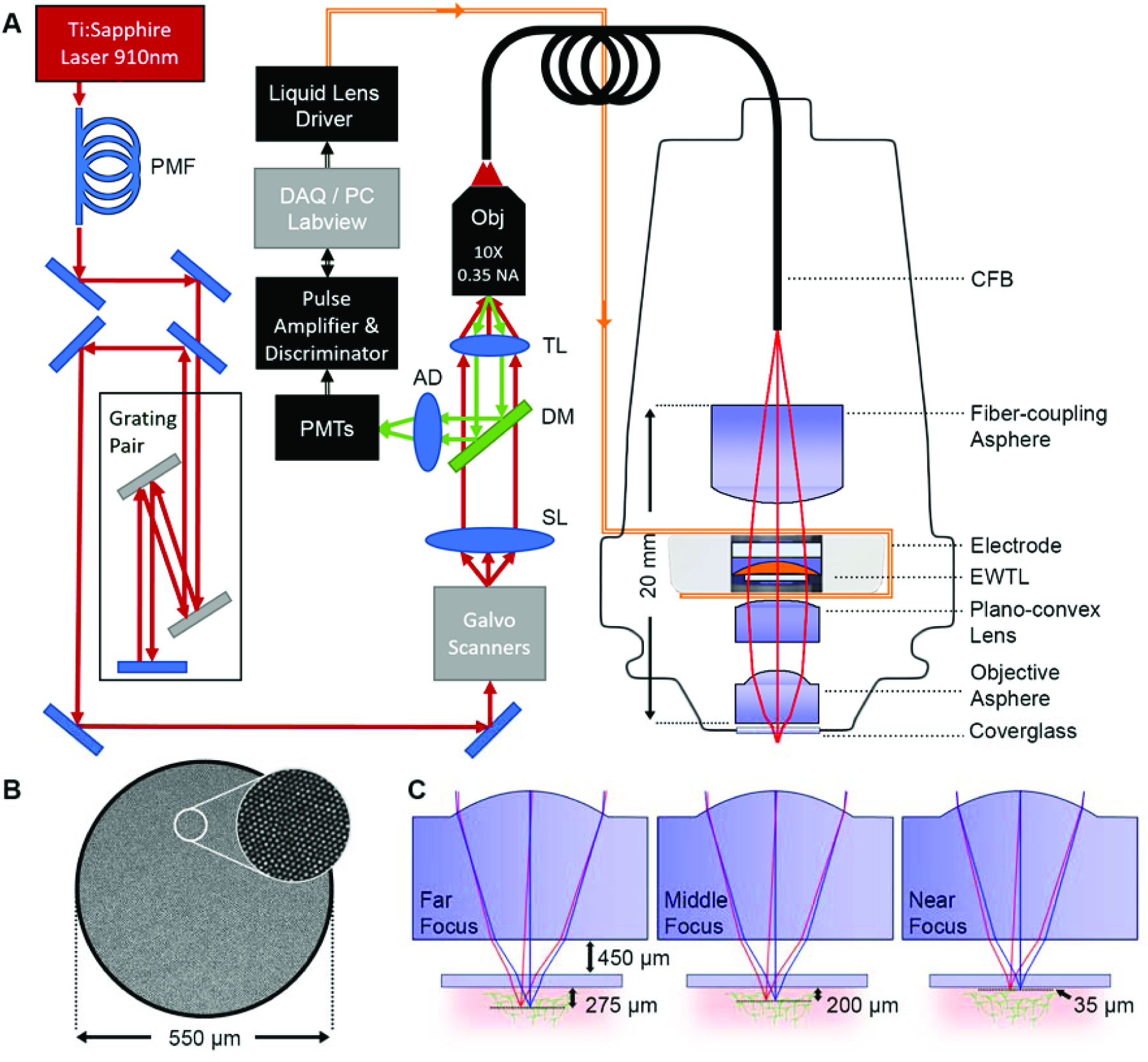
2P-FCM imaging system. **A.** Pulses from a Ti:Sapphire laser at 910 nm are spectrally broadened through a polarization maintaining fiber (PMF) and pre-chirped using a grating pair pulse stretcher. The excitation laser is scanned across the surface the coherent fiber bundle (CFB) through a galvanometric scanning mirror pair, scan lens (SL) and tube lens (TL) relay, and focused with a 10X 0.35 NA Olympus objective. Distal to the CFB, the 2P-FCM optics focus the excitation light from the CFB cores onto the sample through an electrowetting tunable lens (EWTL). Fluorescence emission is collected by the 2P-FCM optics, relayed through the CFB and directed to a photon-counting photo-multiplier tube (PMT) after passing through a dichroic mirror (DM), achromatic doublet (AD), and filter. The PMT signal is amplified and transformed to logic levels to be detected by the data acquisition system on the computer (DAQ/PC). The computer also controls the voltage applied to the EWTL through a driver and flexible electrode. **B.** Detail of the fiber-core pattern of the CFB. C. Zemax simulation of 2P-FCM axial-scanning using specified EWTL lens parameters from manufacturer. The working distance to a #1 coverslip (thickness ∼140 µm) is 450 µm.

The miniature imaging optics for the 2P-FCM are shown enlarged on the right side of **Figure 1A**. The optical system was designed in Zemax optical design software. The 910 nm excitation light from the output of the CFB is collimated through a fiber-coupling asphere (83-710, Edmund Optics), passed through the EWTL to alter the FL, and focused onto the tissue through a plano-convex lens (49-177, Edmund Optics) and short FL asphere (355151, Thorlabs). The emitted fluorescence from the sample is collected through the same optics and coupled back into the CFB. Since the imaging system is designed for accurately focusing the 910 nm light, chromatic mismatch causes the green light collected back through the optics to be defocused at the CFB surface. Fortunately, the large aperture of the CFB means that even heavily defocused fluorescence emission light, such as that scattered through deep tissue is collected. The nominal magnification of the imaging system, from the CFB to the tissue, is 0.4X and the numerical aperture (NA) is 0.45.

The EWTL operates through the principle of electrowetting, in which two immiscible fluids with different refractive indices are influenced by an applied electric field that adjusts the curvature of the interface and thus the FL of the tunable EWTL^27^. The EWTL in this design is a commercially available lens (Arctic 316, Varioptic) with a clear aperture of 2.4 mm diameter and outer diameter of 7.8 mm. Tuning the FL of the EWTL is accomplished by applying an AC voltage of between 25 and 60 V_RMS_ at 1 kHz from a proprietary lens-driver board and via a flexible electrode that is clamped on either side of the lens casing. The maximum specified optical power range of this EWTL is -16 to +36 diopters, although it can be driven at voltages up to 70 V_RMS_ to achieve slightly shorter FL. The EWTL is highly resistant to any motion or vibration as the surface tension forces dominate when the density of the two liquids is matched^28^.

The imaging system is designed to minimize magnification change, maximize the axial-scanning range, and maximize the working distance of the 2P-FCM. **Figure 1C** illustrates the predicted working distance of the 2P-FCM at three EWTL focus settings, in which the working distance is 450 µm from a ∼150 µm thick glass window and the axial-scanning range beyond the window is predicted to span 35 to 275 µm. The predicted imaging properties from Zemax at these three settings are summarized in **Table 1**. A small change in magnification of ∼ 5% through the axial-scanning range was tolerated in order to reduce the total length of the miniature optics.

**Table 1.**
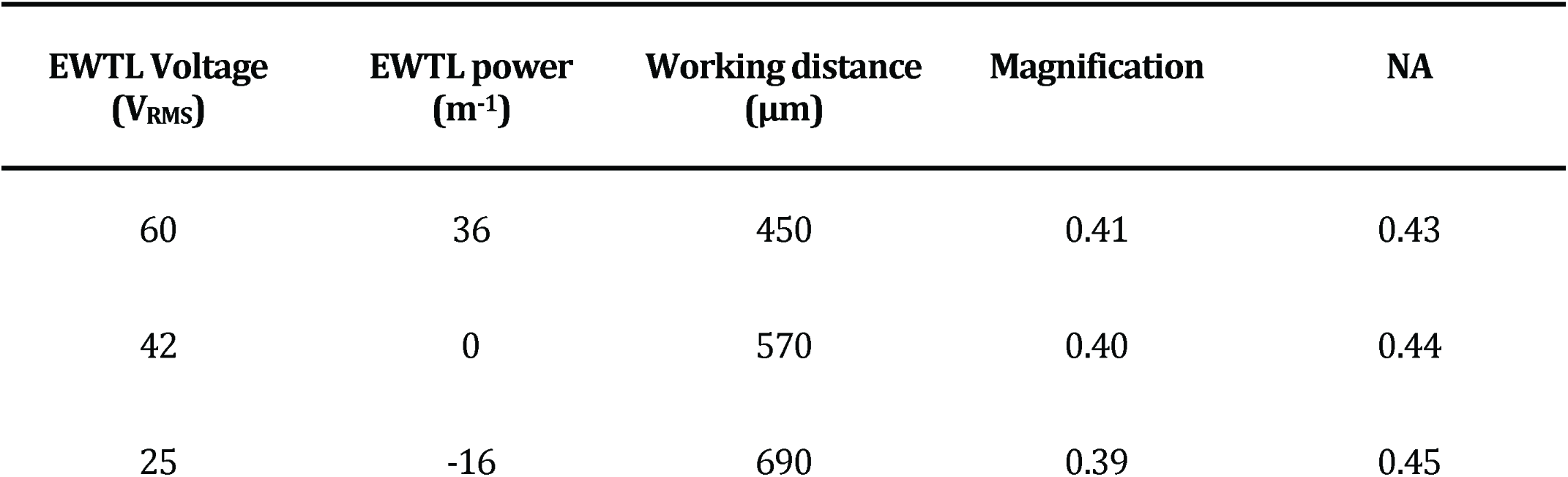
2P-FCM Zemax model optical parameters for different EWTL settings

### Image resolution and axial-scanning range

We characterized the lateral and axial resolution and the axial-scanning range of the 2P-FCM by imaging 2 µm diameter yellow-green fluorescent micro-beads embedded in clear agarose. The lateral resolution is limited by the average spacing of the fiber cores in the CFB. Using a calibrated objective lens, we measured the inter-core spacing to be ∼4.5 µm, which agrees with previously published results^29^. With our magnification factor of ∼0.4x, the theoretical lateral sampling at the object is ∼1.8 µm.

The micro-bead samples were imaged with the 2P-FCM at sequential focal planes by tuning the EWTL focus in discrete steps to obtain a Z-stack. The images were processed to remove the fiber-pixelation pattern as described in the *Methods*. **Figure 2A** shows side-projections of beads as imaged by a 20x 0.75 NA objective (left) and the 2P-FCM (top-right). The same region of the agarose-bead sample was imaged in both cases, such that the majority of the same beads appear in both Z-stacks. This made it possible to compare directly their apparent size and axial location to determine the desired parameters. The beads appear larger than the predicted demagnified fiber-core spacing because each bead was sampled by multiple cores. The coupling to multiple cores is apparent in unprocessed Z-projections shown in **Figure 2B**, which correspond to the substacks indicated in **Figure 2A**.

**Figure 2.**
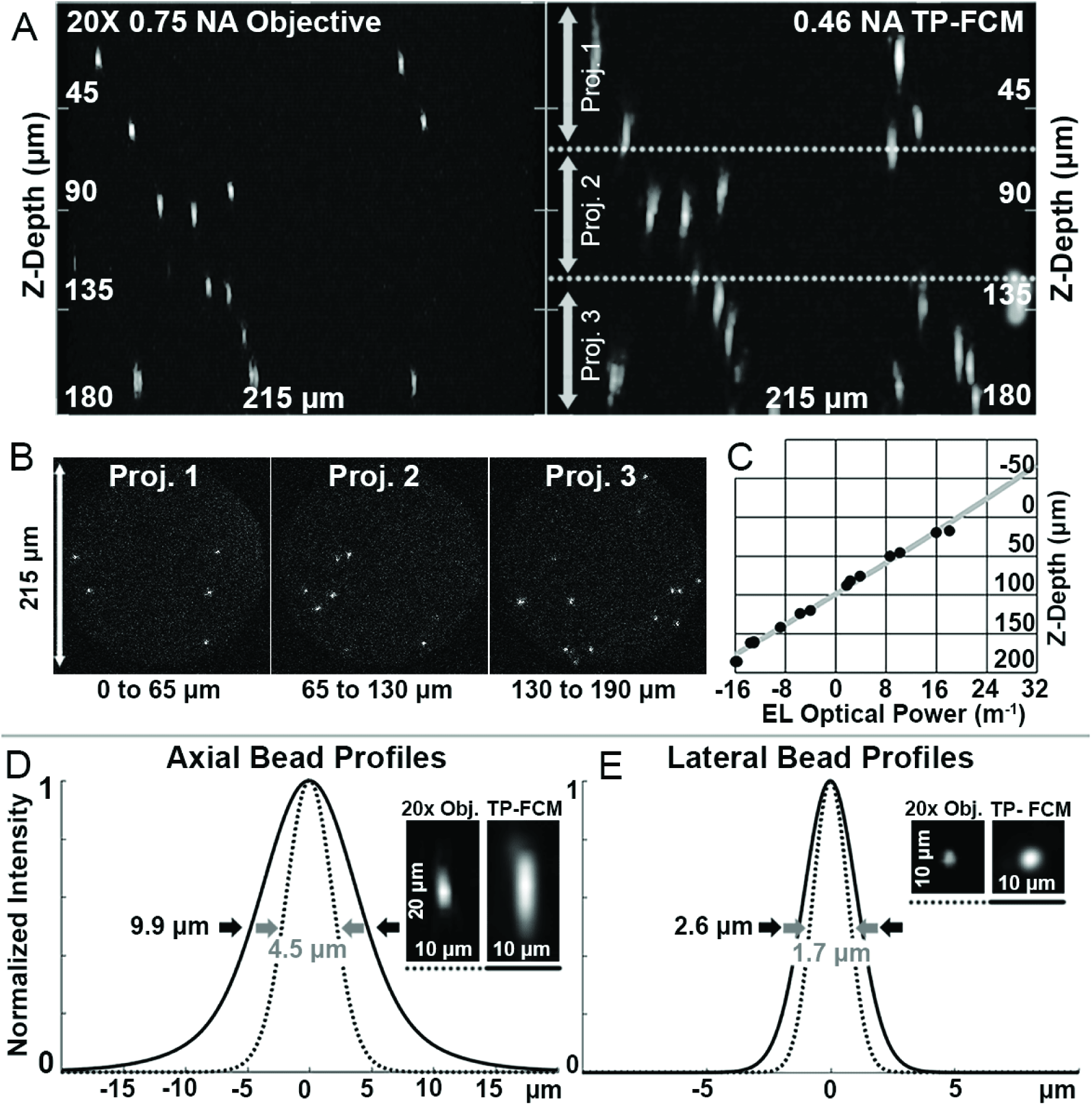
Resolution and testing of axial-scanning range. **A.** Left: Side-projection of ∼2 µm diameter beads suspended in clear agarose and imaged with a 20x 0.75 NA Olympus objective using 910 nm excitation light. Axial focusing was accomplished by a motorized stage. Right: Same volume imaged with 2P-FCM at 910 nm. Axial focusing was accomplished by tuning the focus of the EWTL. **B.** Sub-stack projections of 2P-FCM bead images from regions indicated in A (right panel). **C.** Grey line is the predicted scan range as the EWTL optical power is changed, as modeled in Zemax. Black circles show measured positions of individual beads determined using a motorized stage. **D.** Axial and lateral profiles of beads acquired with 2P-FCM (solid lines) and 20x objective with motorized stage (dashed lines). Profiles are Gaussian fits to averages of 5 beads. Insets show processed images of beads.

The predicated focal plane through the EWTL focusing range from Zemax and actual bead positions are shown in **Figure 2C** (grey line and black dots, respectively). By extrapolating the linear fit of the bead locations to the extents of the scan, we recover that the full axial-scanning range of the 2P-FCM is ∼180 µm from 60V_RMS_ to 25V_RMS_. The axial position versus input voltage is linear, so the axial-scanning depth per volt can be estimated as 5 µm/V. The full axial-scanning range did not span the range predicted (240 µm predicted vs. 180 µm measured) due to under-performance by the EWTL at the high-end of the optical power range. We were able to rule out optical alignment issues by analyzing the limits of possible implementation errors using Zemax. These analyses indicated that any physical alignment error or static lens discrepencies could not account for the change in axial-scanning range. The focal range of the EWTL was tested by measuring the distance from the lens to the focus spot when imaging a distant light source. The optical power at the high voltage setting of 60 V_RMS_ was measured to be ∼ 30 m^-1^, lower than the specified 36 m^-1^ and is the cause of the shorter axial-scanning range.

The lateral resolution was measured after processing the bead images with the interpolation method described in the *Methods* section to remove the fiber-core pixelation. The average lateral and axial line profiles of 5 beads measured from different focus positions were fit to a Gaussian function. **Figure 2D** shows the fitted axial profiles measured using a 20x 0.75 NA microscope objective to be 4.5 µm FWHM (dotted grey line) and with the 2P-FCM to be 9.9 µm FWHM (solid black line). **Figure 2E** shows the fit of the lateral profiles for the 20x objective to be 1.7 µm FWHM (dotted grey line) and with the 2P-FCM to be 2.6 µm (solid black line).

The lateral bead size is larger than diffraction limit as it is limited by the fiber bundle spacing. With a bead size of ∼2 µm, non-uniform sampling of the bead with multiple fiber cores causes a larger effective lateral profile. The axial profile measurements of both the 20x objective at 0.75 NA and the 2P-FCM at 0.45 NA are similar to what is expected from the diffraction-limited calculations.

### Three-dimensional volume imaging

As a first demonstration of the 2P-FCM with axial-scanning, we imaged a thick slice (∼1 mm) of fixed mouse cortical tissue with GFP-labeled oligodendrocytes. Axial-scanning was accomplished by sequentially tuning the EWTL across the full focal range in 36 steps of 1 V_RMS_ per step, or 5 µm per step amounting to a total axial-scanning range of 180 µm. The images were processed to remove the fiber-core pattern as described in the *Methods* section. A 3D volume image was created using ImageJ/Fiji software and is shown in **Figure 3A**. We observe over 200 individual cells in this volume. **Figure 3B** shows a single imaging plane after processing to remove the fiber-core pattern through interpolation, which shows the cell bodies of the oligodendrocytes clearly but is unable to resolve the finer processes. This demonstration shows that we are able to visualize deep into scattering tissues to resolve a large number of cells.

**Figure 3.**
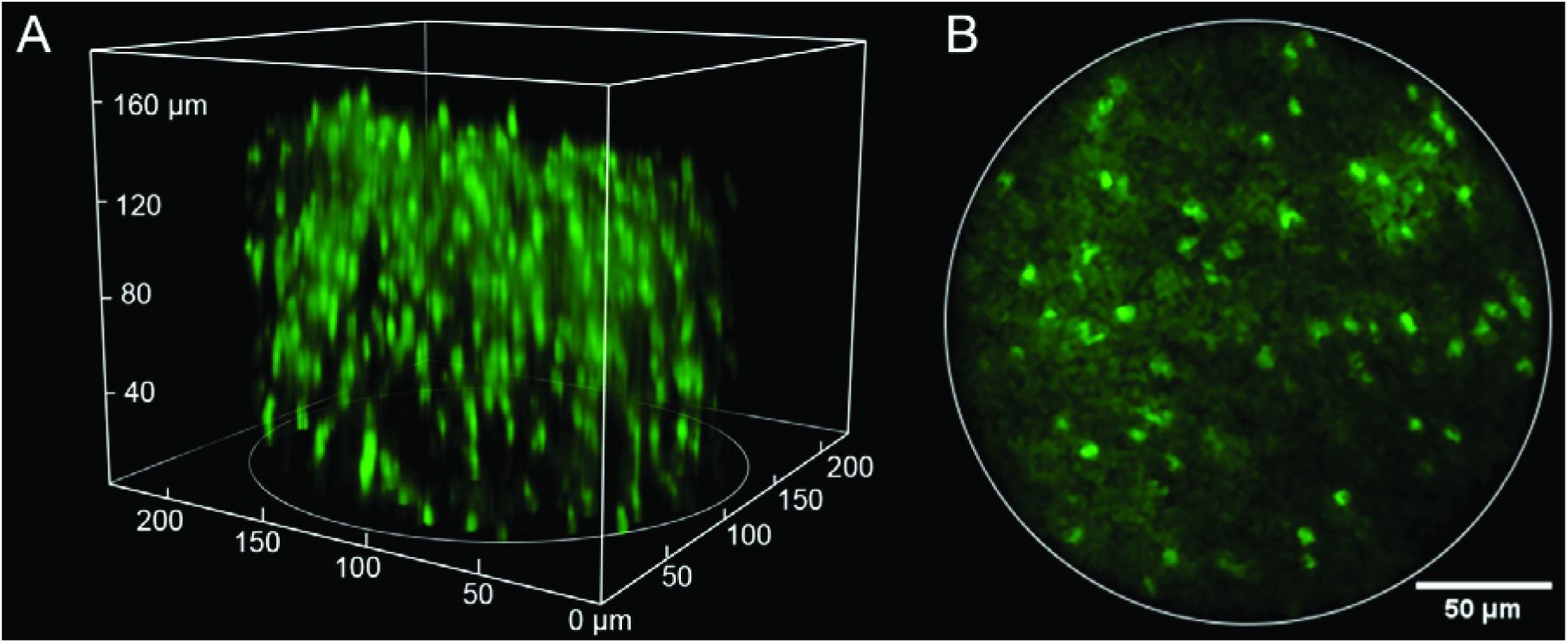
**A.** Two photon image showing a 3D volume acquired by the 2P-FCM (220 × 220 μm lateral x 180 μm axial) with over 200 cells in the image. The sample is fixed mouse cortical tissue (4% paraformaldehyde) expressing green fluorescent protein (GFP) driven by proteolipid protein promoter. The GFP labels oligodendrocyte cell bodies in the cortex. **B.** Processed image of a single slice in the stack after filtering to remove pixelation pattern.

### Tilted-field imaging tests

Tilted-field imaging increases the functionality by imaging over different regions of interest in three dimensions. Because the active axial-scanning is done with the EWTL, it does not require any mechanical motion or moving parts, making the scans very repeatable and stable. The tilted-field scan is accomplished by driving the EWTL with a control signal that varies across the scan field. **Figure 4** shows the results of performing this test on fixed, thick brain tissue with sparse fluorescence signal from neurons expressing GCaMP6s. 25 total angles were acquired, 12 in each direction, varying the voltage on the EWTL from 40 to 50 V_RMS_ across the slow-axis of the raster scan. The tilted image fields are shown sequentially in **Supplementary Movie 1**. The ability to perform scanning at different angles allows coverage of the brain region of interest for a given experiment. In general, the waveform applied to the EWTL was a simple sawtooth function, however new methods for voltage-function shaping can optimize the speed and dynamics of tunable lenses based on electrowetting^30^.

**Figure 4.**
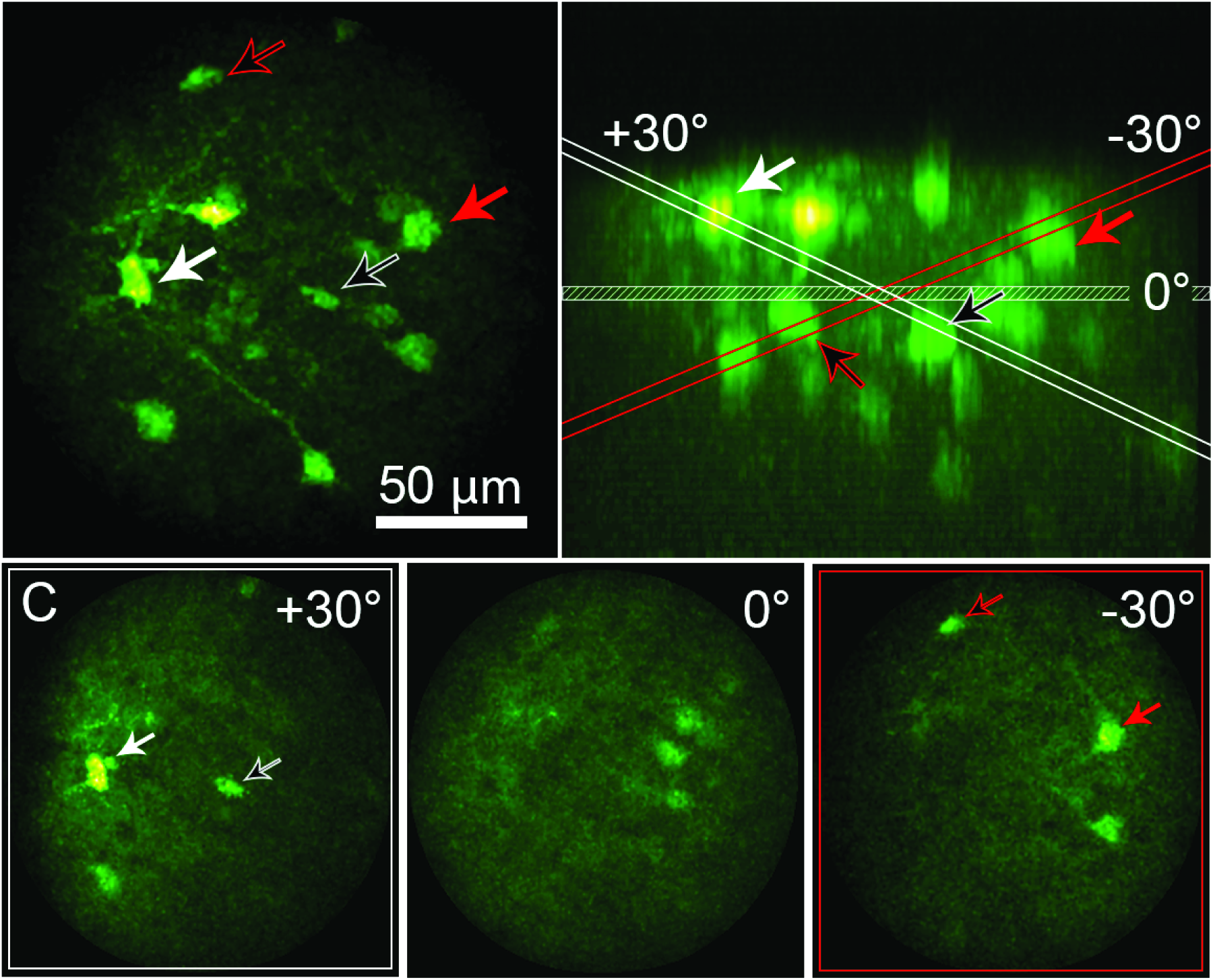
Tilted-field scan enabled by rapid focusing of the EWTL lens. **A.** Maximum intensity projection of a thick coronal section of mouse brain, expressing GCaMP6s in neurons, acquired by the 2P-FCM. Arrows indicate cell bodies that are visible during tilted-field scanning. **B.** Side (XZ) projection of the image volume in A. The same cell bodies are indicated by the arrows. The planes for the horizontal and angled scan limits are indicated and color coded red for the -30 degrees scan, and white for the +30 degrees scan. **C.** Images showing the tilted-field scans, indicating the same cell bodies that are shown to intersect with the red or white planes in B.

### Two-color imaging

Multi-color imaging an important feature for researchers who want to investigate the co-localization or relative abundance of certain physiological markers. The 2P-FCM offers an advantage in that two-photon excitation at a single wavelength can excite a number of different fluorophores with separated emission spectra that can be imaged simultaneously. A common example is GFP and tdTomato, which are both efficiently excited around 900 nm with 2PE, but have spectrally separate emission bands. **Figure 5** shows two-color imaging with 900 nm 2PE of tdTomato and GFP simultaneously with a 20x Olympus objective and separately with the 2P-FCM. The two images were acquired using the same detectors and filters. The 2P-FCM therefore can provide many of the flexible features of a 2PE microscope.

**Figure 5.**
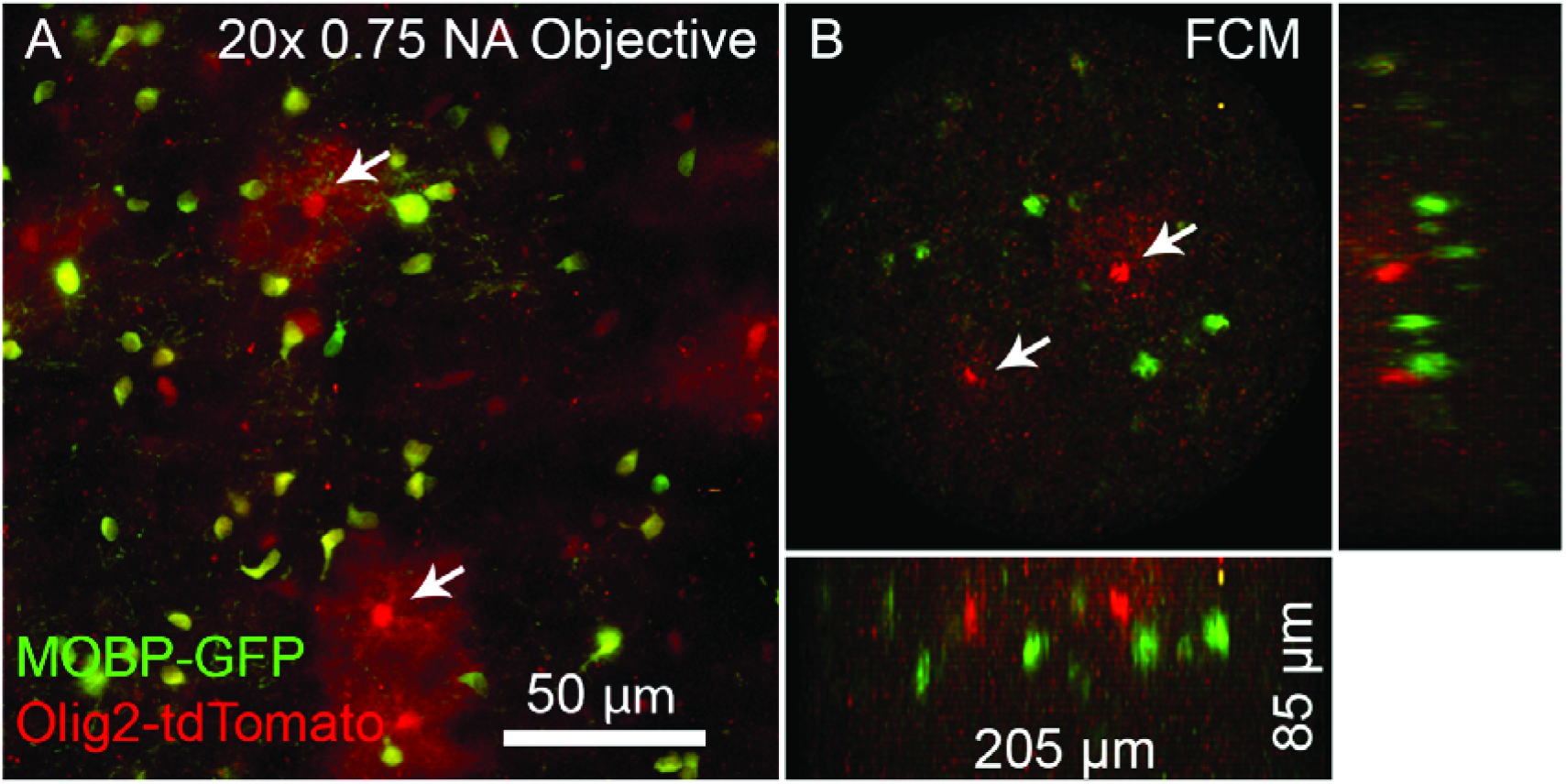
Two-color imaging of GFP-expressing mature oligodendrocytes and tdTomato-expressing oligodendrocyte lineage cells and sparsely labeled astrocytes (*Mobp-EGFP; Olig2-Cre; R26-lsl-tdTomato* triple transgenic mice). **A.** Maximum intensity projection of a region of brain tissue acquired with a 20x 0.75 NA objective. Yellow cells are oligodendrocytes (Green and Red), while red-only cells are astrocytes and oligodentrocytes cells. Two astrocytes are marked with arrows, and are easily identified by the characteristic bushy morphology and two are marked with arrows. **B.** The same tissue imaged with the 2P-FCM, using the same detectors, filters, and excitation wavelength. Green and red cells are visible in the field, with likely astrocytes marked by arrows.

### Stability of 2P-FCM for longitudinal animal imaging

For freely-moving mouse imaging, the 2P-FCM package should be light-weight, resistant to motion and vibration, and yield good repeatability for imaging over long time spans. We tested the optical and mechanical aspects of the 2P-FCM by imaging neurons expressing GCaMP6s in the somatsensory cortex of a freely-moving mouse. For each imaging session, the 2P-FCM was attached to the baseplate excerting minimal torque on the skull (**Figure 6A**). **Figure 6B** shows a photo of a mouse with the 2P-FCM attached on top of an existing head-fixation bar for head-fixed 2PE imaging, allowing both methods to be used with the same mouse. The CFB and electrode wire for the EWTL were draped passively over a horizontal post to guide them into the cage. By itself, the assembled 2P-FCM with the CFB weighs 2.5 g. When attached, the 2P-FCM, CFB, and electrode added only ∼3 g to the mouse’s weight. The mouse was able to move freely in an 8” by 12” area. The movement was somewhat restricted by the length (1.0 m) and torsional resistance of the CFB, preventing rotation of more than ∼360^°^. We found that after short acclimation (∼30 minutes) mice were able to traverse the entire region and habituated to restrictions on rotation. After a few sessions mice could perform normal behaviors, such as walking on a rodent exercise wheel **(Supplementary Movie 2)**. Future implementation may include a commutator with a rotation encoder for realignment in post-processing^12^.

**Figure 6.**
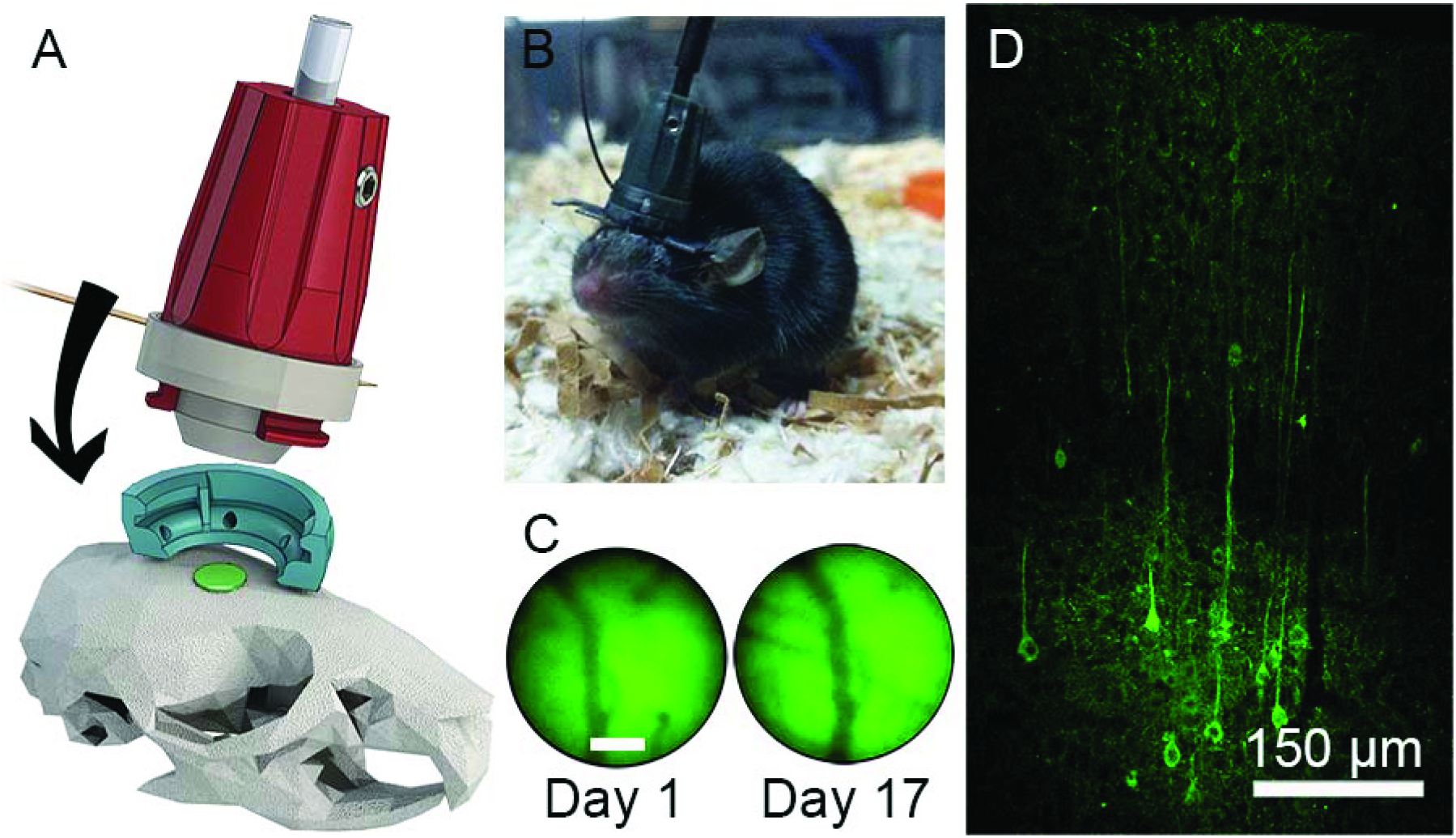
Mouse implant and verification of GCaMP6s expression. **A.** Attachment mechanism of 2P-FCM onto baseplate attached to the mouse skull. **B.** Photo of a behaving mouse with 2P-FCM attached. **C.** Verification of stability of imaging FOV over 17 days, showing the same blood vessels with epifluorescence measured through the FCM. **D.** Confocal image of the neocortex for a fixed cortical coronal slice of the mouse’s brain 12 weeks post-injection of AAV5 expressing GCaMP6s driven under control of the synapsin promoter.

**Figure 6C** shows a wide-field green epifluorescence image taken through the 2P-FCM under blue light illumination. The left image shows background fluorescence and vasculature on the first day of 2P-FCM attachment. The right image shows the same field, indicating that the baseplate was stable over 17 days with only a slight lateral shift in alignment. The expression level in neurons virally transfected with GCaMP6s in the cortex was verified by histology. **Figure 6D** shows a 20 μm-thick confocal stack of a coronal section of a perfused mouse 12 weeks post injection. Neuronal cell bodies are seen in layers 4/5, as well as layers 2/3 in lower density, where we performed imaging with the 2P-FCM.

### *In vivo* mouse imaging during behavior

We used the 2P-FCM to perform *in vivo* 2PE imaging of neuronal activity in the motor cortex of an awake and mobile mouse expressing GCaMP6s and tdTomato in cortical neurons. **Figure 7A** shows a photo of the selected FOV for imaging because of the low density of vasculature, taken through one of the eyepieces of the microscope. The head-adapter portion was attached to the mouse skull with acrylic cement to maintain repeatable imaging over the same FOV. Before the awake-imaging sessions, the mouse was very briefly anesthetized (< 30 seconds) for quick attachment of the 2P-FCM to the head-adapter.

**Figure 7.**
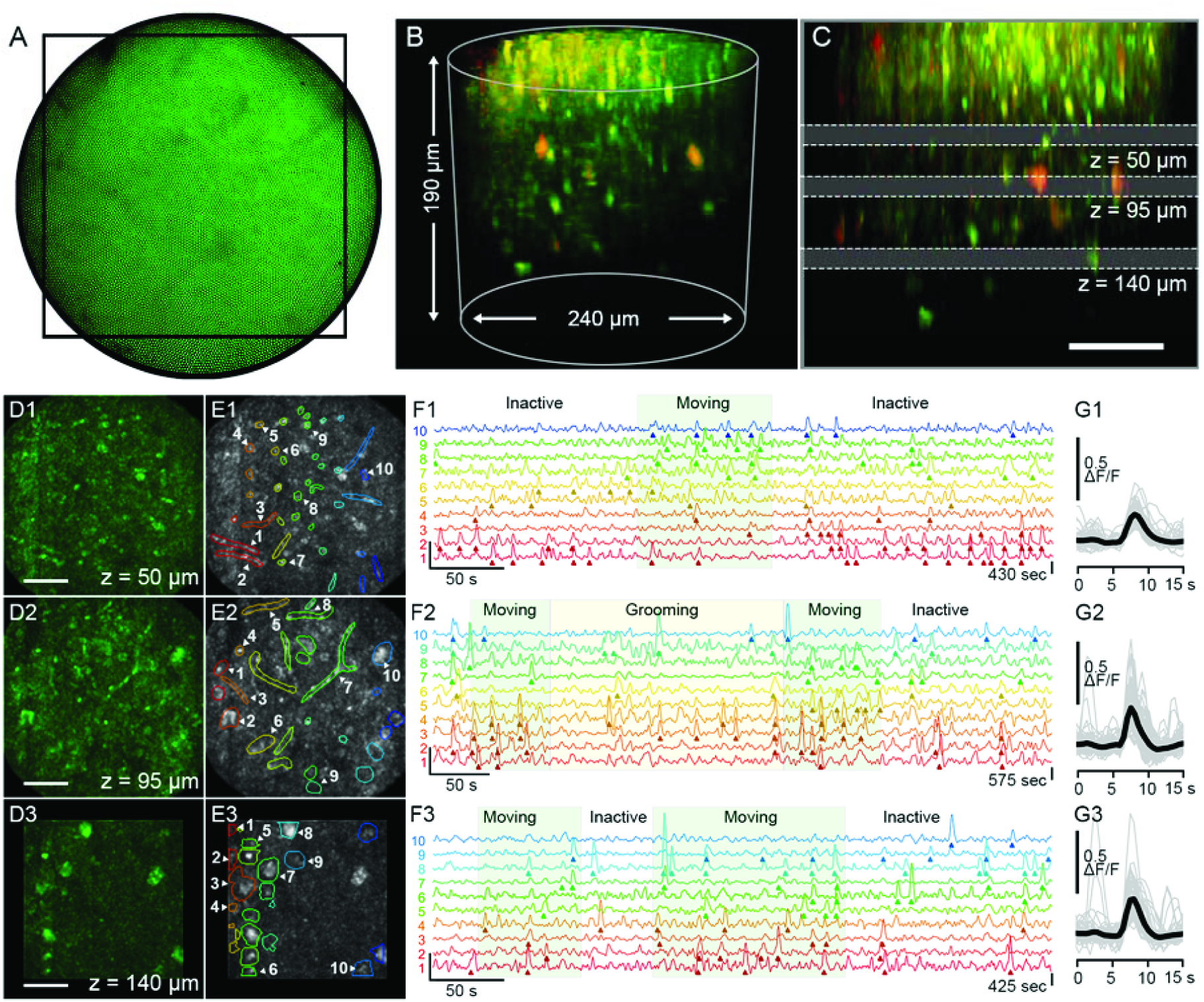
Two-photon Ca^2+^-imaging in an awake and freely-moving mouse using the 2P-FCM. A. Widefield camera image of the epifluorescence taken through the FCM showing showing vasculature in the FOV. **A** black rectangle indicates the acquisition region for the following images. **B.** 3D projection of a Z-stack acquired by sequentially imaging and focusing through the tissue using the EWTL. **C.** Side projection of the same Z-stack indicating the three depths at which time series were acquired: at 50 µm, 95 µm, and 140 µm depth. **D.** Maximum intensity projections of image frames that coincide with Ca^2+^-transients, showing structural changes through the focal depths. Scale bar is 50 µm. **E.** Selected ROIs that contain fluorescence transients that exceed the 6 SD threshold. Time traces for ten representative ROIs are selected for each depth. **F.** Detailed time-courses of the ΔF/F signal for the ten indicated regions. Regions of mouse behavioral activity from the camera recording are overlayed on the traces. **G.** All identified transients, aligned by the peak ΔF/F signal, shown in gray with averaged intensity signal in black.

**Figure 7B** is a 3D-projection of a Z-stack taken by tuning the EWTL focus from 62V to 25V at 0.5V per step (∼190 µm at 2.5 µm per step). The signal from GCaMP6s is shown in green and the signal from the more sparsely labeled tdTomato is shown in red. Near the surface of the tissue are small, bright structures that are likely the neuronal processes in the neuropil. Starting at approximately 60 µm depth is a region of mixed processes and cell bodies, in which shadows of vasculature can be seen. At approximately 120 µm below the surface, there is an increase in the density of cell bodies, indicating the beginning of neocortical layers 2/3. (**Supplementary Movie 3**). **Figure 7C** shows a side projection of the same volume with annotations indicating the three depths chosen for activity recordings, one placed in each of the three described sub-regions. Temporal activity recordings were performed at depths of 50 µm, 95 µm, and 140 µm (EWTL settings: 52, 43, and 34 V_RMS_). For the 50 and 95 µm depths, the rectangular FOV was 210 × 210 µm, indicated by the box in **Figure 7A**. At the 140 µm depth, a smaller FOV of 140 × 140 µm was used to account for increased frame time needed for imaging at greater depth. The frame acquisition rate varied from 1.3 to 2.5 Hz based on the optimal dwell time per pixel needed to see cellular details. Higher frame rates can be easily achieved by reducing the FOV.

The results of the recordings for each depth are summarized in **Figure 7D-F**, separated into three time-series taken at the different axial focus depths. Only the GCaMP6s channel was used for these analyses. For each time-series, an intensity projection following the interpolation-based image processing, is shown in **Figure 7D**. In **Figure 7E** the same projection is shown with a sub-set of manually selected regions-of-interests (ROIs) that have been shown to contain at least one fluorescence transient that exceeds the threshold of 6 times larger than the standard deviation (SD) of the background intensity fluctuations (see *Methods* for details). Near the surface, at 50 µm, the regions of activity are much smaller or are elongated, indicating that the activity is likely from small axonal or dendritic processes in the neuropil, which are often perpendicular to the imaging plane. At the deepest 140 µm depth, the active regions are round objects 10-20 µm in diameter, which are likely cell-bodies. At the middle depth of 95 µm, there is a mixture of processes and cell bodies. Five ROIs for each depth are selected and the ΔF/F traces are shown in **Figure 7F**, with detected fluorescence transients indicated by arrows. The general behavior of the mouse, determined by video taken by a camera above the cage, is overlayed on the plot. During the “Moving” periods, the mouse is traveling around the behavioral cage. During the “Grooming” periods the mouse is moving on the camera but is not traveling around the cage. In the “Inactive” periods the mouse is not moving appreciably. A quantification of the transient timing and behavioral motion is shown in **Supplementary Figure 1A**. The time-series taken at the 50 µm and 140 µm depths correlate significantly with behavioral motion (see **Supplementary Figure 1B**). The data taken at the 95 µm depth does not correlate significantly with behavioral motion, due to a large number of fluorescence events occuring during the “Inactive” period. It is possible that smaller but meaningful behaviors, such as sniffing or whisking, were missed by the camera and video-analysis. Interestingly, some of the data indicate a selective responsiveness of the ROIs to behavior, which is especially noticeable at the 50 µm depth traces. In Figure 7G, the detected transients from these ROIs are aligned by the time point corresponding to the rising edge of the transient, overlayed by the averaged transient shape. The shape follows the expected exponential shape of a neuronal Ca^2+^-transient detected via GCaMP6s, with a distribution of durations between 2 and 4 seconds (see **Supplementary Figure 1C**)^31^. We also recorded activity while the mouse recovered from anesthesia, resulting in a gradual buildup of neuronal activity (see **Supplementary Figure 2** and **Supplementary Movie 4**). Lateral motion artifacts were present during some of the recordings. We used a motion correction algorithm to correct for the laterally shifting field (see *Methods*). The results of the motion correction indicate a time-averaged motion artifact magnitude of < 2 µm, but had peaks up to 10 µm in some cases, (see **Supplementary Figure 1A**). The intensity of bright cells that did not exhibit fluorescence activity did not vary with the motion artifacts, so we conclude that the extent of the motion in the axial-dimension is lower than the depth-of-focus for the 2P-FCM (<10 µm). Overall, these motion artifacts were similar to those reported in head-fixed 2PE imaging studies^32^ and wide-field head-attached imaging^9^. The 2P-FCM did not become loose or dislodged during the imaging sessions, which lasted between 1 to 4 hours.

## Discussion

We demonstrated the first head-mounted 2P-FCM with active axial-scanning to attain 3D-imaging of neuronal activity in freely moving mice. The optical design was optimized for resolving neuronal somata and some neuronal processes with a lateral resolution of 1.8 µm and axial resolution of 10 µm while also allowing for a large FOV of 240 μm diameter and axial-scanning range of 180 μm depth. The 2P-FCM recorded neuronal activity at three structurally distinct depths of a freely-moving mouse. The design of the compact attachable head mount was stable over weeks, demonstrating the ability for longitudinal studies.

Head-fixed 2PE microscopy is useful because of the ability to image cortical neuronal and dendritic activity in awake animals^2^. Importantly, recent advances have allowed mesoscopic measurements in large numbers of neurons^33^. Some aspects of free movement can be simulated by placing the animal in a virtual reality situation^34^. However, restriction of head movement limits neuronal imaging to a subset of behaviors. Our 2P-FCM will be useful to study neuronal activity during behaviors that involve the movement of the animal such as social behaviors^7^, prey capture^8^, fine motor coordination and learning^35^ and odorant plume tracking^36^. As a proof-of-concept, we demonstrate correlation of motor-cortex neuronal activity with unguided movement of a freely-moving mouse. A full validation of the capability of the 2P-FCM to record neuronal activity correlated to complex behavior will be undertaken in future studies.

Important for these experiments is an imaging system that is resistant to motion and vibration. Motion artifacts were minimized by the rigid and compact design and use of the EWTL as the only active focusing element attached to the animal. The 2P-FCM is light weight and applies ∼3 g of vertical force on the mouse, but tethering the mouse to the CFB does restrict rotational motion. In our experiments, the mice were able to acclimate to the restriction and were able to navigate the entire behavioral field as well as perform more complex tasks, such as walking on an exercise wheel. In the future, incorporation of a commutator will allow more freedom of rotation for the animal^12^.

The 2P-FCM design also provides a great degree of flexiblity for the user. The CFB permits the selection of arbitrary ROIs within the FOV, which can be increased in size for slower and larger region scanning, or decreased in size for faster and smaller regions when targeting small clusters or single cells. The active axial-scanning afforded by the EWTL allows live and rapid readjusment of the focal plane without needing to interfere with the behavior. This is essential for focal alignment with the restricted depth-of-focus of 2PE microscopy, and also allows for rapid re-selection of focal planes that can be programmed for sequential or triggered acquisition over the course of a long experiment. Compared with other methods of axial-scanning in miniature microscopes, the EWTL is simple in implementation, small in size and weight, and immune to orientation and vibration. The EWTL is also capable of more rapid variation of the focus rapid 3D imaging. We demonstrated more complex 3D scans with tilted field imaging, which may be important for accessing features that are not confined to a single focal plane. In this study we altered the EWTL FL slowly through the scan, but the EWTL has been shown to be capable of varying the FL ∼50 Hz or greater when driven at resonance^37,38^. In the future we can implement more efficient driving functions for the EWTL to achieve high-speed axial scanning, opening the potential for random-access scanning for truly volumetric data collection^3,4,39^.

Compared to a bench-top 2PE laser-scanning microscope system, there are some limitations with the 2P-FCM. The 3D imaging volume (240 μm diameter x 180 μm depth) is smaller than the volume imaged by head-fixed 2PE imaging. The lateral resolution is limited by the CFB core-spacing, which reduces the resolution below that achievable with comparable NA objectives, and the imaging quality is reduced by the spacing and non-uniform 2PE sensitivity of the fiber-cores. We demonstrate that the 2P-FCM resolution is sufficient to record from cell bodies and some processes even at > 100 µm depth while also providing access to many more behavioral options than head-fixed imaging. Post-processing with intensity masking and interpolation is able to effectively reduce the intensity variation and remove the honeycomb pixelation pattern that afflicts CFB 2PE imaging. This opens up options for temporal analyses that would not have been possible on the raw fiber-image, and can be used to extract more spatial information that may unveil subcellular dynamics as well^40^. The choice of the CFB as the imaging relay also means that the entire optical system can be placed in-line, since the EWTL itself is a transmissive element and the CFB can be used for relaying a proximally scanned excitation spot as well as acting as a large aperture for fluorescence collection from the entire FOV.

The 2P-FCM with active axial-scanning is a mechanically and optically simple solution to neuronal imaging of freely-moving mice. In the future our 2P-FCM can be paired with GRIN lenses to image deep brain regions^41^ and can be used for simultaneous optogenetic stimulation^42,43^.

## Methods

### Microscope setup

A home-built 2PE laser scanning microscope was used to perform imaging through the 2P-FCM. The excitation source is a Spectra-Phyics MaiTai HP Deepsee ultrafast laser, with ∼80 fs pulses tuned to a center wavelength of 910 nm and operating at 80 MHz repetition rate. The beam power is controlled by a half-wave plate on a motorized rotation mount (Conex-AG-PR100P, Newport) followed by a Glan-Taylor polarizer (GT10-B, Thorlabs). The 1 mm beam is coupled into a 0.5 m length PMF (PM780-HP, Thorlabs), with APC termination to reduce back-reflections. The output of the PMF is collimated with a fixed fiber-collimation lens (F220APC-780, Thorlabs). The 1.0 mm beam is then expanded through a 3.75x beam expander to reduce the average intensity on a grating-pair pulse stretcher, with lenses: -40 mm FL (ACN25-040-B, Thorlabs) and 150 mm FL (LB1437-B, Thorlabs). The gratings are reflective ruled gold with a density of 300 grooves/mm (49-572, Edmund Optics) separated by 265 mm. The grating-pair stretcher is used to apply ∼66,000 fs^2^ of negative GDD to the laser pulse to compensate for the positive dispersion from the 0.5 m length PMF, 1.0 m length CFB (FIGH-15-600N, Fujikura), and additional chirp resulting from non-linear self-phase modulation (SPM) in the CFB at the power levels used in the experiments. The precise grating separation distance was determined empirically by maximizing the fluorescence signal while imaging a fluorescent test slide (Chroma Technology) through the 2P-FCM. After traveling through the grating-pair, the beam size is reduced by 2.5x with a reverse beam expander with lenses: 100 mm FL (LB1676-B, Thorlabs) and -40 mm FL (ACN25-040-B, Thorlabs). The beam is scanned through a galvanometric mirror scanning system (6215H, Cambridge Technologies). The beam is relayed through a 50 mm FL scan lens and 180 mm FL tube lens in an Olympus IX71 microscope. The beam is focused and laterally scanned on the surface of the CFB through a 10X 0.35 NA Olympus UPLANSApo objective lens. A XYZ-translation stage (CXYZ05, Thorlabs) is used to accurately focus and align the fiber to the focus of the objective lens. The beam propagates through the CFB and then through the distal miniature optics onto the tissue, resulting in fluorescence excitation. The fluorescence emission is collected through the same optical path and the CFB. Because the 2P-FCM optics are not chromatically corrected, green emission light is projected onto a ∼50 µm diameter area on the CFB surface. We measured 532 nm light to be transmitted at ∼67% efficiency through the CFB when imaged through multiple fiber-cores. The objective collects the fluorescence emission from all fiber-cores of the CFB, which is separated from the excitation path by a primary dichroic mirror (T670LPXR, Chroma Technology). A second dichroic mirror (FF562-Di02, Semrock) splits the red and green fluorescence for two-color detection. Each color line of the fluorescence emission is focused onto a large-area photon counting photo-multiplier tube (PMT) (H7422P-40, Hamamatsu) with an achromatic doublet lens (LB1761-A, Thorlabs). The output pulses from the PMTs are amplified by a high-bandwidth amplifier (ACA-4-35db, Becker & Hickl GmbH) and are converted to logic-level pulses by a timing discriminator (6915, Phillips Scientific). The pulses are counted by a data-acquisition card (PCIe-6259, National Instruments) at a rate of 20 MHz. The counts are sampled and binned by pixels and converted into an image in custom software in Labview (National Instruments) that also controls the EWTL driver.

### Ultrafast laser-pulse propagation through fiber-bundle

Chromatic group delay dispersion (GDD) of the 1.0 meter length glass fiber cores of the CFB results in broadening of the pulse-width which can significantly reduce the two-photon excitation fluorescence signal. The GDD can be pre-compensated with a grating-pair pulse stretcher to add negative chirp to the pulse. In the linear regime, this will result in a transform limited pulse out of the CFB. However, propagation of high peak-power pulsed light through a small core fiber results non-linear self-phase modulation (SPM) that affects the spectral properties of the pulse. In the special case where the pulse has a large negative chirp, SPM in the fiber causes spectral narrowing and results in temporal broadening of the pulse out of the CFB. High peak-power pulse propagation through optical fibers for multiphoton imaging has been explored extensively in previous work^26,44-46^. An effective solution is to first induce spectral broadening by propagating the transform-limited pulse directly out of the laser in a PMF before applying negative chirp from the grating pair. As the pulse propagates through the CFB, the spectrum narrows but is balanced to obtain the initial spectral width from the laser, and thus a similar pulse duration.

In this work, we used a 0.5 m length PMF to spectrally broaden the pulse before using the diffraction grating-based pulse stretcher to apply negative GDD to the pulse before launching into the CFB cores^24,26^. We measured the resulting pulses through the CFB using frequency resolved optical gating (FROG) to recover the intensity and phase information of the resulting electric field. With only the grating-pair and no spectral pre-broadening, the original pulse width of 80 fs FWHM with spectral width of 16 nm FWHM, (**Supplementary Figure 3A**), is stretched to a temporal width of 640 fs before propagating through the CFB. At laser powers below ∼ 2 mW, the pulse is re-compressed back to 80 fs temporal width with a 13 nm spectral width after propagating through a single fiber core of the 1.0 m-long CFB (**Supplementary Figure 3B**). When the laser power is increased to 40 mW out of the CFB, the pulse bandwidth is narrowed to 5.4 nm through SPM and the temporal pulse-width is broadened to 635 fs (**Supplementary Figure 3C**). Finally, when the PMF fiber is used for pre-broadening the spectrum, the output pulse at 60 mW is recompressed with a temporal width of 110 fs and a spectral width of 16 nm (**Supplementary Figure 3D**). The measurement of the phase indicates third order dispersion (TOD), which grating-based pulse stretchers do not compensate. A shorter pulse width can be achieved by using a compression system that also provides negative TOD, such as a grism pair, as previously described^44,47-49^.

High-peak power pulses incident on the proximal glass surface of the CFB also result in unwanted background resulting from autofluorescence and inelastic raman scattering^50^. We find that the background signal from the fiber surface is greatly reduced when the pulse is pre-chirped, resulting in fewer non-linear interactions at the proximal fiber surface.

### 2P-FCM mechanics and attachment

The enclosure for the 2P-FCM optics is designed in Solidworks 3D CAD software (Dassault Systèmes). The packaging is split into three sections: top, bottom, and baseplate, shown in **Figure 6A**. The top section contains the fiber bundle ferrule, held in place by two set-screws, and the fiber-collimating asphere. The bottom-section contains the objective lenses. The unmounted lenses are held in by friction in precisely sized openings. The top-section has two curved tabs that interface with slots in the bottom section, which help to ensure reproducable alignment. The EWTL and the electrode are sandwiched between the bottom section and the top section with an O-ring that ensures good electrical contact. The flat-flex electrode cable exits the enclosure through a small slot between the sections. The top-section tabs have a single thread at the end, which interfaces with the baseplate as shown in **Figure 6A**. In this way, the basepate is pulled up against the bottom-section by the top-section and greatly improves rigidity when attached to a moving animal. The baseplate is designed with ridges and holes to improve adhesion of the cement for attachment to the animal skull. The entire enclosure is 3D-printed using a high-resolution projection-based resin printer (Kudo3D Titan 1), with a resolution of 50 μm. 3D printing allows optimization of the prototype and easily enables design changes, such as the inclusion of GRIN lenses for deep brain imaging^51^.

The CFB and EWTL electrode may change the effective weight on the mouse when attached. We measured the mouse in its cage before and after attaching the 2P-FCM. The added weight to the mouse with the 2P-FCM attached varied from 2.0 g to 3.0 g as the mouse traveled around the cage, indicating that the CFB sometimes relieves some of the weight of the CFB and sometimes increases it. The CFB has a minimum bending radius of ∼60 mm and is relatively stiff, resulting in an upwards force on the attached 2P-FCM that counters its weight. Also, the CFB cannot be reached by the mouse, preventing any distractions or damage to the CFB or electrode during the behavioral experiments.

### Bead sample preparation

Resolution and axial scan range measurements were performed by imaging fluorescent beads embedded in agarose (A9414, Sigma-Aldrich) and a USAF 1951 resolution target (38-257, Edmund Optics). 2 µm yellow-green fluorescent beads (F8853, Invitrogen) were used to measure axial scanning extent as well as lateral and axial resolution.

Low melting-point agarose was prepared at a concentration of 0.5 % in water. The 2 μm diameter fluorescent yellow-green beads were diluted in the agarose to a concentration of ∼2.0 × 10^7^ beads/mL. Approximately 2.0 mL of solution was placed on a #1 coverglass and allowed to set at room temperature. The beads were imaged in sequential axial planes by a 20x 0.75 NA Olympus UPLANSApo objective with a motorized stage and separately by the 2P-FCM by changing the voltage applied to the EWTL.

### Mouse neuronal imaging

All experiments were approved and conducted in accordance with the Institutional Animal Care and Use Committee of the University of Colorado Anschutz Medical Campus. Male 3-month old C57BL/6 mice were anesthetized by intraperitoneal ketamine-xylazine injection. The skin above the target site was numbed by lidocaine injection and retracted to expose the skull. The mouse was injected with a mixture of adenoassociated virus (AAV) similar to procedures in Ref. ^5^. The AAVs used in the mixture expressed GCaMP6s under the synapsin promoter (AAV5.Syn.GCaMP6s.WPRE.SV40, UPenn Vectorcore) and tdTomato under the synapsin promoter (AAV5-eSYN-TdTomato, Vector Biolabs). These AAVs were injected with a ratio of 2:1 respectively with a total injection volume of 1.2 μL delivered with a glass micro-pipette through a 0.5 mm hole drilled at the target site. The coordinates of the injection targeted the motor cortex, 0.4 mm anterior to bregma and 1.2 mm lateral to the midline, at a depth of 300 μm^52^.

One month after injection the mice were implanted with an optical cranial window near the injection site, using standard techniques as previously described^53^. Briefly, mice were anesthetized by isoflurane inhalation and the skin under the scalp was numbed by subcutaneous lidocaine injection. The skin above the skull was removed to expose the injection site and skull surface. A 2 mm square window of skull was removed immediately anterior to the injection hole to expose the dura mater. The opening was covered with a 2 mm square #1 coverglass and secured in place with cyanoacrylate glue. Dental acrylic cement (C&B-Metabond) was used to cover the skull surface. The presence of fluorescence signal was confirmed with standard 2PE microscopy using a 20x 1.0 NA Zeiss Plan-Apochromat water-immersion objective.

The baseplate attachment procedure is similar to what has been described for other miniature head-attached microscopes^54^. While the mouse was still anesthetized, the 2P-FCM was held with a micromanipulator (Sutter MP-285) and positioned above the cranial window until fluorescent signal could be observed using widefield epifluorescence through the 2P-FCM. A final FOV was chosen by searching for good cellular expression by performing 2PE imaging through the 2P-FCM. The baseplate was then secured to the existing acrylic around the cranial window with black acrylic cement (Jet Acrylic, Lang Dental). After allowing to set for ∼30 minutes, the 2P-FCM was removed, leaving the baseplate in place, and the mouse was allowed to recover.

For imaging, the mouse was briefly anesthetized with isoflurane inhalation. The baseplate was carefully gripped by thumb and forefinger and the 2P-FCM was inserted and secured with a quarter-turn. The EWTL electrode was connected to light-gauge wires draped, along with the CFB, over a horizontal metal post above the behavior cage. After recovery, the mouse was allowed to wander freely in an 8” by 12” plastic cage that was illuminated by a small battery powered lamp. The light was placed carefully to avoid illuminating any of the 2PE microscope detection optics, which would cause excess background signal. The external light caused a slight increase in the background signal of about 15% in the green channel. We also tested red illumination, which is not visible to the mouse but allows for imaging with the camera^55^. Red illumination did not result in an increase in background signal in the green channel. An overhead webcam (Logitech C615) recorded the mouse behavior during the imaging experiments.

Imaging was performed at 910 nm without modulating the output power with depth since we were able to run the system at near full-power without causing saturation of the PMTs. The time-averaged power out of the Ti:Sapphire oscillator was 1040 mW, while the time-averaged power measured after the 2P-FCM was 60 mW. The majority of the power loss in the system is due to the grating pair (44% transmission) and the low NA air objective lens on the 2P microscope (65% transmission). The transmission can be improved with more efficient gratings and use of an objective lens optimized for near infrared transmission.

### Image processing

The images from the 2P-FCM show a honeycomb pixelation pattern due to the packing of the cores of the CFB. Several methods have been described to depixelate images from CFBs^56-58^. The simplest methods involve low-pass filtering with either a blurring function^24^ or masking the image in the frequency domain^59^. However, two-photon imaging through a CFB has the additional complication of amplifying the non-uniformity of the fiber cores. Each core is assumed to have a unique sensitivity, due to the variability in diameter, shape, NA, and amount of cladding between adjacent cores. This manifests as discrete variations in image intensity across the FOV.

To address this, we created a flat-map of the full CFB fiber-cores by imaging a fluorescent test slide (Chroma Technologies) with the 2P-FCM. The flat-map stores the centroid coordinates of the cores and their corresponding sensitivity. The processing was performed with custom software in Matlab (Mathworks). Each image to be analyzed was registered to the flat-map, which allowed identification of the cores. The relative sensitivity of each core was compensated by dividing by the flat map values. The honeycomb pattern was eliminated by using the natural nearest neighbor interpolation method^60^. A Savitsky-Golay filter was used to reduce the added single-pixel noise introduced by the core multiplication factor during flat-normalization. The most recent version of the fiber-processing code is made available on Github^61^.

For processing of temporal scans, each frame was processed with the same flat-map alignment so that the cores are static in the field. Once the honeycomb pattern and CFB-induced intensity variation was removed, a clustering algorithm was used to identify regions of interest (ROI) of high-correlation^62,63^. Significant changes in cytosolic Ca^2+^ were identified as changes in fluorescence larger than 3 standard deviations above baseline within each ROI.

## Acknowledgments

We gratefully acknowledge Robert Cormack for fruitful discussions. We also acknowledge Nicole Arevalo for assistance with animal care, Elizabeth Gould and Wendy Macklin for providing brain tissue from Plp-EGFP mice. The study was funded by National Science Foundation DBI-1353757, CBET-1631912, IIP-1602128 and National Institutes of Health NS099577, DC000566, NS048154. Some of the imaging in this study was performed in the University of Colorado Anschutz Medical Campus Advance Light Microscopy Core that is supported in part by Rocky Mountain Neurological Disorders Core Grant Number P30NS048154 and NIH/NCATS Colorado CTSI Grant Number UL1 TR001082.

## Author Contributions

B.N.O. was involved in project conception, experimental design, performed experiments, data processing/analysis, and wrote the manuscript. G.L.F. contributed to experimental design and performed experiments. M.M. performed experiments. V.M.B and J.T.G were involved in project conception. E.J.H. performed experiments and contributed with expertise in cortical imaging. D.R. was involved in project conception, data analysis and manuscript writing. E.A.G. was involved in project conception, experimental design and direction, data analysis and manuscript writing. All authors were involved in editing the manuscript.

