## Supplementary Materials for "Three dimensional two-photon imaging of neuronal activity in freely moving mice using a miniature fiber coupled microscope with active axial-scanning"

**Supplementary Material**

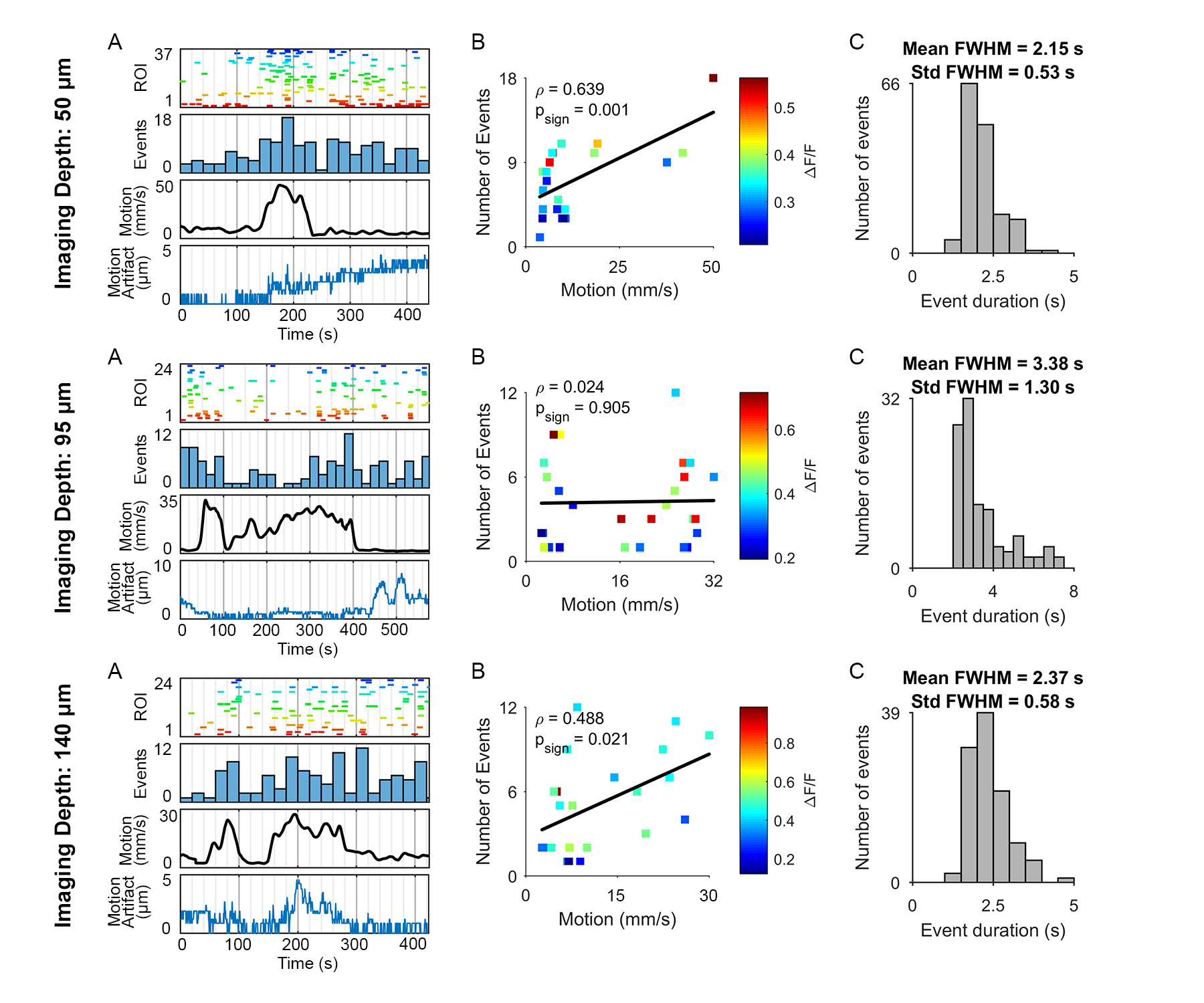

**Supplementary Figure 1.** Behaving mouse calcium-imaging data from three imaging depths (50 µm, 95 µm, 140 µm) acquired using the 2P-FCM. **A.** Summary of neuronal and behavioral activity through imaging duration. Top: Raster plot of fluorescence event times for all active ROIs, coded by color. Top-middle: Histogram of the number of transients that occur within each 20 second window. Bottom-middle: Optical flow analysis of the mouse movements during the imaging study, showing the average velocity of movement during each time window. Bottom: Magnitude of motion correction required to stabilize images. **B.** Correlation analysis of average behavioral motion velocity against the number of fluorescence events that occur in each 20 second window. Each event is color-coded by peak ΔF/F. The black line is a linear regression of all events. The Pearson’s correlation coefficient (ρ) and p-value for the correlation are indicated. **C.** Histogram of FWHM for fluorescence events.

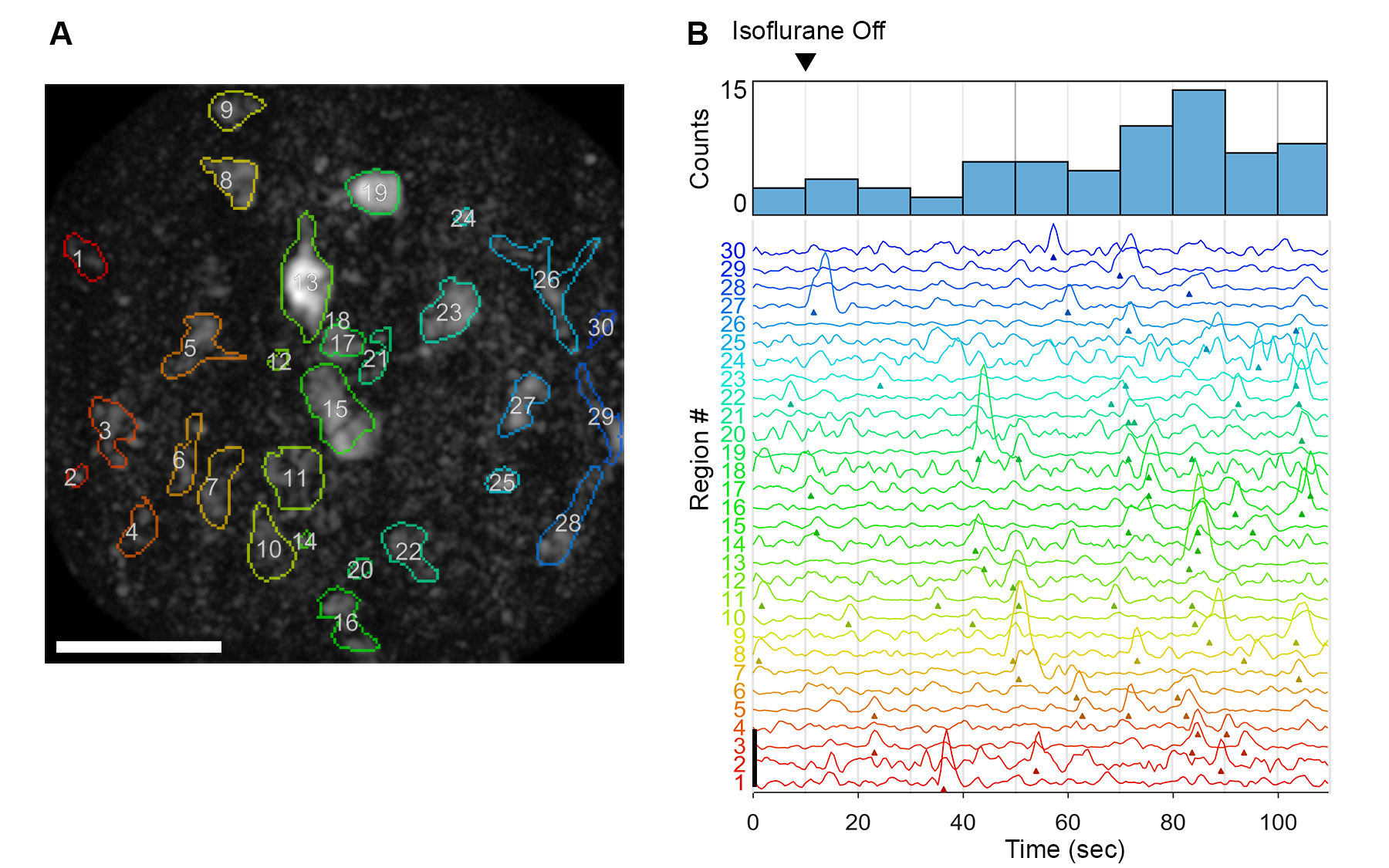

**Supplementary Figure 2.** Recovery of neuronal activity after discontinuing isoflurane gas anesthesia. Focus was set to a depth of approximately 100 µm into the cortex. Images were acquired at 2 Hz for 110 seconds, with the isoflurane gas delivery being discontinued after 10 seconds of imaging. **A.** Maximum intensity projection of time series overlayed with colored manually drawn ROIs. Scalebar is 50 µm. **B**. Recorded events above the 6 standard deviation threshold showing an increase in neuronal activity following removal of anesthetic gas. Top: Events histogram with 10 second bins. Bottom: ΔF/F signals from each ROI, with arrows signifying beginning of detected event. Scale line in bottom left indicates ΔF/F of 1.5.

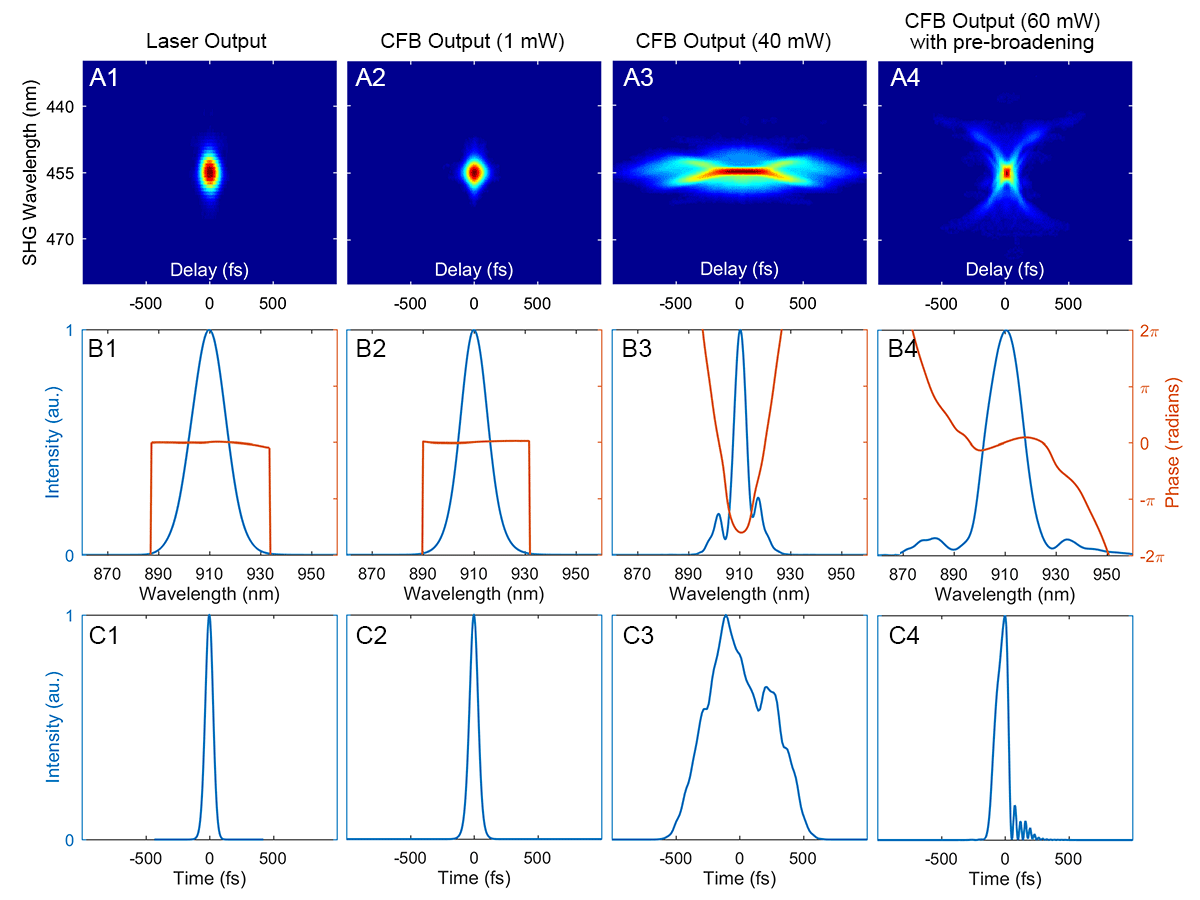

**Supplementary Figure 3 -** Measurement of ultrashort pulse propagation through 1.0 m-long CFB by FROG technique at 910 nm. **A.** Spectrogram of SHG FROG. **B.** Retrieved spectral intensity (blue) and phase (orange). **C.** Retrieved temporal intensity profile. Intensity units are normalized. **A-C. (1)** Measurement taken directly from laser output. Temporal pulse width is 72 fs FWHM and spectral width is 16 nm FWHM. **(2)** Measurement taken after pulse is propagated through the grating-pair pulse stretcher and a single-core of the CFB at low power (power measured at output of CFB is ~ 1 mW). Temporal pulse width is 80 fs FWHM and spectral width is 12.6 nm FWHM. **(3)** Same grating pair stretcher setting as in (2) but at higher power (power measured at output of the CFB is ~ 40 mW). At high power, temporal pulse width is increased due to spectral narrowing caused by self-phase modulation of the negatively stretched pulse and non-linear dispersion. Temporal pulse width is 635 fs FWHM and spectral width is 5.6 nm FWHM. **(4)** Measurement taken after pulse is propagated through the PM fiber before the grating-pair to pre-broaden the pulse to compensate for the spectral narrowing. The increased spectral width and increased grating-pair separation result in a short output pulse. Temporal pulse width is 110 fs FWHM and spectral width is 16 nm FWHM.
